# Dynamic actin crosslinking governs the cytoplasm’s transition to fluid-like behavior

**DOI:** 10.1101/774935

**Authors:** Loïc Chaubet, Abdullah R. Chaudhary, Hossein K. Heris, Allen J. Ehrlicher, Adam G. Hendricks

## Abstract

Cells precisely control their mechanical properties to organize and differentiate into tissues. The architecture and connectivity of cytoskeletal filaments changes in response to mechanical and biochemical cues, allowing the cell to rapidly tune its mechanics from highly-crosslinked, elastic networks to weakly-crosslinked viscous networks. While the role of actin crosslinking in controlling actin network mechanics is well-characterized in purified actin networks, its mechanical role in the cytoplasm of living cells remains unknown. Here, we probe the frequency-dependent intracellular viscoelastic properties of living cells using multifrequency excitation and *in situ* optical trap calibration. At long timescales in the intracellular environment, we observe that the cytoskeleton becomes fluid-like. The mechanics are well-captured by a model in which actin filaments are dynamically connected by a single dominant crosslinker. A disease-causing point mutation (K255E) of the actin crosslinker α-actinin 4 (ACTN4) causes its binding kinetics to be insensitive to tension. Under normal conditions, the viscoelastic properties of wild type (WT) and K255E+/- cells are similar. However, when tension is reduced through myosin II inhibition, WT cells relax 3x faster to the fluid-like regime while K255E+/- cells are not affected. These results indicate that dynamic actin crosslinking enables the cytoplasm to flow at long timescales.

## Introduction

The actin cytoskeleton is a network of filaments crosslinked by actin binding proteins. The cytoskeleton provides mechanical support and drives cell motility and morphological changes^1^. Crosslinking proteins undergo continuous cycles of binding and unbinding, enabling the cell to act as an elastic solid at short timescales and as a viscous fluid at long timescales^2,3^. To characterize the mechanics of the active, viscoelastic cellular environment, we developed methods to probe cytoplasmic mechanics in living cells over timescales ranging from 0.02 to 500 Hz using an optical trap (OT). Active microrheology as applied here is insensitive to non-equilibrium active cellular processes like motor protein-based transport and cytoskeletal dynamics because only the response that is coherent with the applied displacement is analyzed^4,5^. OTs can apply a local deformation directly in the cytoplasm of living cells, while other commonly used active methods such as atomic force microscopy (AFM) and magnetic twisting cytometry apply a local deformation at the cell surface. The viscoelastic response is described by the frequency-dependent complex shear modulus G*(f) = G’(f) + iG’’(f) where the real part G’(f) and the imaginary part G’’(f) correspond to the elastic (storage) and viscous (loss) modulus, respectively. While the magnitudes of G’(f) and G’’(f) reported in the literature vary greatly between different measurements, their frequency dependence is well conserved across different cell types and experimental conditions^4^. G’(f) and G’’(f) increase with frequency following a weak power law G*(f) ∼ f^α^ with α = 0.05-0.35 from ∼1 Hz to ∼100 Hz^4-18^, and a stronger power law G*(f) ∼ f^β^ with β = 0.5-1.0 for frequencies above ∼100 Hz^4,7,13,19,20^. The weak power law has been attributed phenomenologically to the soft glassy material (SGM) theory^12,13,21^, while the strong power law has been explained by entropic filament bending fluctuations^13,22,23^. However, at timescales longer than 1s (i.e. at frequencies below ∼1 Hz) corresponding to cell division, motility and morphological changes, active microrheological data is scarce especially in the cytoplasm of living cells. Microrheology using thermally driven particles is prone to errors at these timescales from the contribution of active processes in the cell^4,5^. In purified actin networks, transient crosslinking by actin-binding proteins governs the viscoelastic properties at long timescales^24-31^. When these networks are deformed at frequencies slower than the crosslinker’s unbinding rate (k_off_), the crosslinkers have enough time to unbind, allowing the filaments to freely slide past one another resulting in a more fluid-like, viscous network (Fig. 1a). At deformation frequencies faster than k_off_, the actin filaments remain highly interconnected resulting in filament bending and a more solid-like, elastic network (Fig. 1b). We hypothesize that this role of dynamic actin crosslinking may allow the actin cytoskeleton to become fluid-like during movement and morphological changes, while also maintaining structural integrity. While active microrheological studies on the cell surface have reported weak power-law scaling of mechanics at timescales down to ∼0.1 Hz^11-18,32^, the distinct mechanical properties in the interior of the cell^4^ at these long timescales remain largely unexplored. OTs are well-suited to measure these mechanical properties and while several studies have reported intracellular viscoelasticity^5,6,8-10,19,20,33^ the mechanics at ∼0.01-1 Hz have not been quantified in detail. Experimental measurements of slow dynamics are challenging due to the long observations required and while *in situ* OT calibration provides more accurate measurements^34-36^, it has often been neglected.

**Figure 1.**
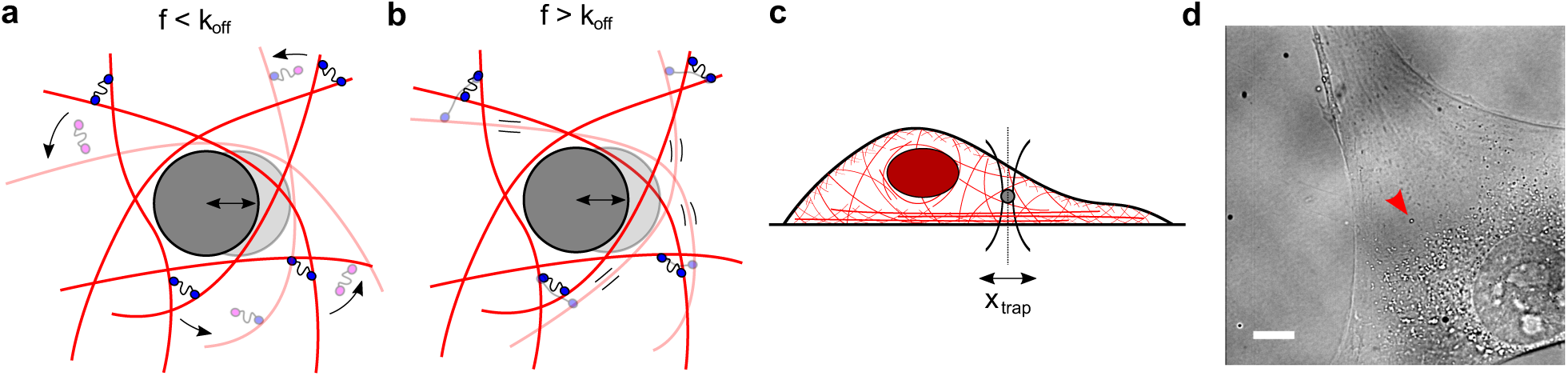
The cytoplasm is a crowded viscoelastic matrix of crosslinked polymers and proteins. **a**, Schematic of the actin network being deformed by a microbead at frequencies (f) slower than the unbinding rate of the crosslinkers (k_off_), before (opaque) and after deformation (transparent). Previously bound crosslinkers (blue heads) detach (magenta heads) and the filaments are free to slide past one another and rearrange (curved arrows), resulting in a more viscous, fluid-like network. **b**, At frequencies faster than k_off_, the crosslinkers stay bound and interlock the filaments, resulting in filament bending and a more solid-like, elastic response. **c**, Schematic of the experimental setup showing the active oscillation of bead in the intracellular environment by the optical trap (x_trap_). Not to scale. **d**, Representative brightfield image of a live cell showing the optically trapped bead (arrow). Scale bar is 10 µm.

## Results and Discussion

To perform simultaneous measurements over a wide frequency range, we optically trap a 500-nm polyethylene glycol (PEG)-coated bead located in the cytoplasm (Fig. 1c,d, Methods) and apply a multifrequency excitation input to the trap position using an acousto-optic deflector (AOD). Because there are significant elastic contributions from the cytoplasm and non-equilibrium active processes, the trap stiffness, k_trap_, and the photodiode constant, β_pd_, cannot be calibrated using traditional methods relying on thermal fluctuations. Here, we fit a simple viscoelastic model to the measured response to calibrate the optical trap *in situ*^35^. The frequency response function (FRF) is obtained from the time domain signal of the photodetector and the trap position (Fig. 2a,b). The magnitude and phase of the FRF, as well as the power spectrum are simultaneously fit to the model to estimate k_trap_ and β_pd_. The magnitude response at frequencies above ∼1 Hz is well-captured by this simple viscoelastic model, with a first inflection point at ∼4 Hz representing the stiffness of the cytoplasm and a second inflection point at ∼200 Hz representing the combined stiffnesses of the cytoplasm and the OT (Fig. 2b, Supplementary Information). The resulting k_trap_ and β_pd_ vary between cells and between experiments due to variable optical properties (Supplementary Fig. 2), reinforcing the necessity for *in situ* calibration. Despite variability among cells, the product of k_trap_ and β_pd_ falls within the same constant range, as reported previously^37^ (Supplementary Fig. 2). The viscoelastic response measured in entangled purified actin networks (Supplementary Fig.3) using this method agrees well with literature^22,38,39,40,41^, and the response measured in a 20% PEG water solution agrees well with data obtained from a rheometer (Supplementary Fig. 4). Once calibrated, k_trap_ and β_pd_ are used, along with the FRF, to obtain the complex shear modulus G*(f)^38^. Combined with multifrequency excitation, this calibration method allows k_trap_, β_pd_ and G*(f) to be obtained in a single continuous measurement, thereby minimizing sources of error (see Supplementary Information). This approach has advantages over other methods used to calibrate the OT in active, viscoelastic media. Methods based on light momentum changes rely on capturing most of the diffracted and scattered light^37^. However, organelles and other structures result in substantial refraction. An alternative method uses the active response to sinusoidal inputs combined with the passive response for *in situ* calibration in viscoelastic medium^19,20,34-36,42-44^. k_trap_ can be obtained from the average k_trap_ values at each input frequency based on the ratio of the real part of the active spectrum to the passive thermal spectrum at the same frequency^19,20,34,36,42-44^. β_pd_ can be calibrated separately using an active oscillation of the stage or the laser combined with the position readings from the camera or photodiode. However, both k_trap_ and β_pd_ are sensitive to sources of error as they are derived from a single frequency point.

**Figure 2.**
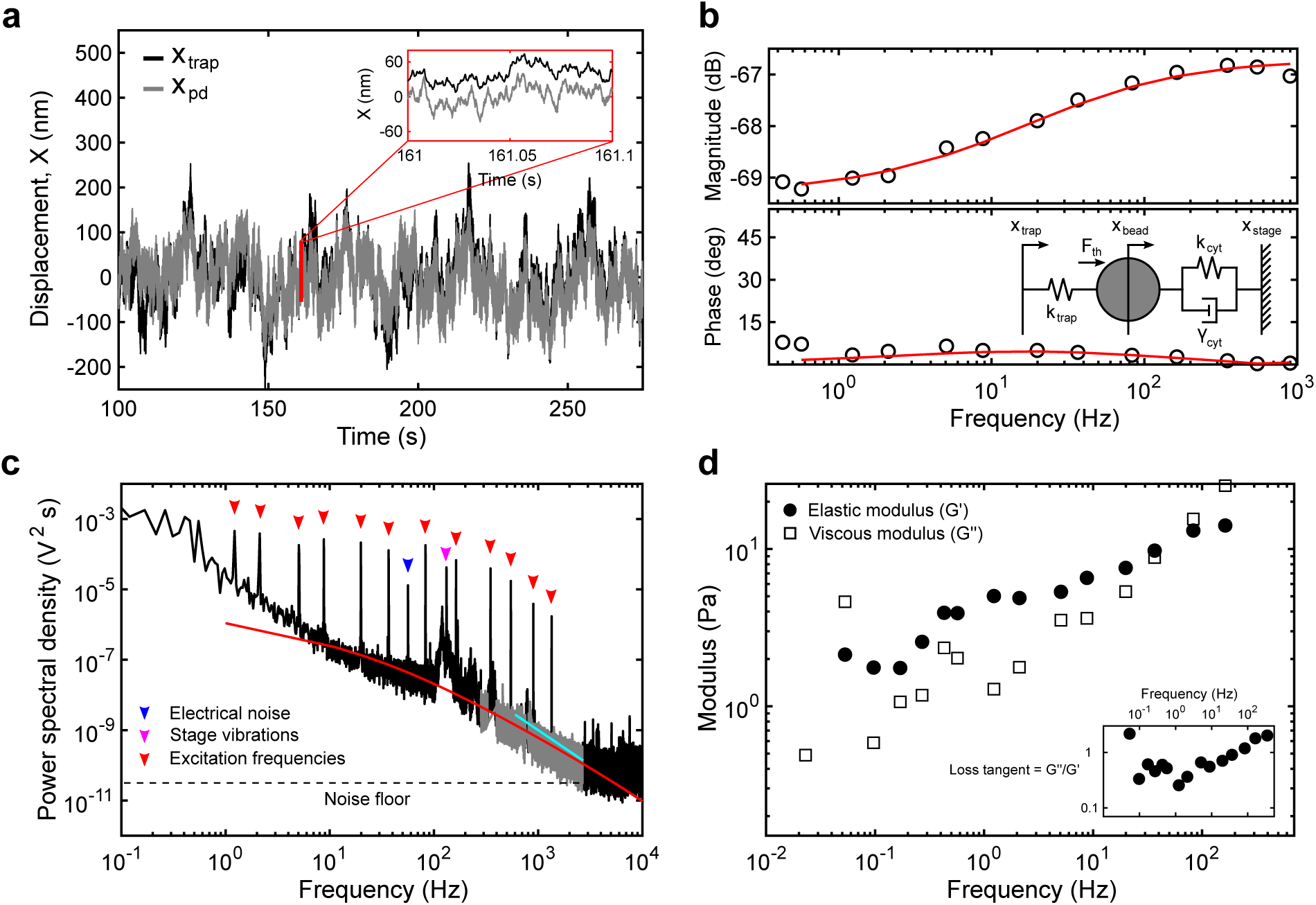
Analytical methods to obtain the viscoelastic properties of the cytoplasm from optical trapping data. **a**, Sample trace of the trap position (x_trap_, black) and photodiode position (x_pd_, gray) in the time domain. Note that the photodiode signal is acquired in volts and converted to nanometers after *in situ* calibration. **b**, The magnitude and phase of the response of the photodiode output signal to movements of the trap at each input frequency is captured by the transfer function. The transfer function above 1 Hz is fit (red) to the active part of a simple viscoelastic model (inset) to obtain the trap stiffness k_trap_ (pN/nm) and photodiode constant β_pd_ (nm/V). The mechanical circuit (inset) depicts the forces experienced by the bead from the optical trap and the viscoelastic environment. **c**, The power spectrum of the output between 250 Hz and 2500 Hz (gray) resulting from thermal fluctuations is simultaneously fit (red) to the passive part of the model. Below 250 Hz, non-equilibrium active cellular processes, electrical noise (blue arrow head) and stage vibrations (magenta arrow head) contribute to the response while above ∼2500 Hz, the noise floor (dashed line) is reached. The active response from the multifrequency excitation (red arrow heads) is visible. A slope smaller than 2 (cyan) indicates constrained diffusion. **d**, The elastic (G’, solid circles) and viscous (G’’, open squares) moduli are obtained for each experiment (see Supplementary Information).

Using multifrequency excitation and *in situ* calibration, we characterize the viscoelastic properties of the cytoplasm of living cells at frequencies ranging from 0.02 to 500 Hz. In WT fibroblasts, the measured moduli can be described by a double power law (Fig. 3a, magenta), yielding a weak power law (α = 0.116 ± 0.021) at frequencies between ∼1 Hz and ∼20 Hz and a stronger power law above ∼20 Hz (β = 0.717 ± 0.062), in excellent agreement with previous observations^5,8,13,19^. However, at frequencies below ∼1 Hz, we observe a pronounced transition to a fluid-like response, apparent in the loss tangent (G’’/G’) which reaches a minimum at ∼1 Hz and increases steadily with decreasing frequencies. This transition to a more fluid-like regime occurs at timescales relevant to slower processes such as cell migration^2,45^ and cell division^3,46^ but is not well captured by the power law model (Fig. 3a, inset). A transition to fluid-like behavior has been observed in mixtures of actin filaments and actin crosslinkers *in vitro*, where the timescale is determined by the crosslinker unbinding kinetics^27-31^. The observation of a similar fluid-like transition in cells is surprising as many crosslinking proteins with different binding kinetics likely contribute to the viscoelastic response. Several possible mechanisms might explain the single dominant timescale for relaxation: (1) the slowest crosslinker dominates the mechanical response, (2) alpha-actinin-4 is the dominant crosslinker in the interior of the cells examined here, or (3) many crosslinking proteins have similar binding kinetics.

**Figure 3.**
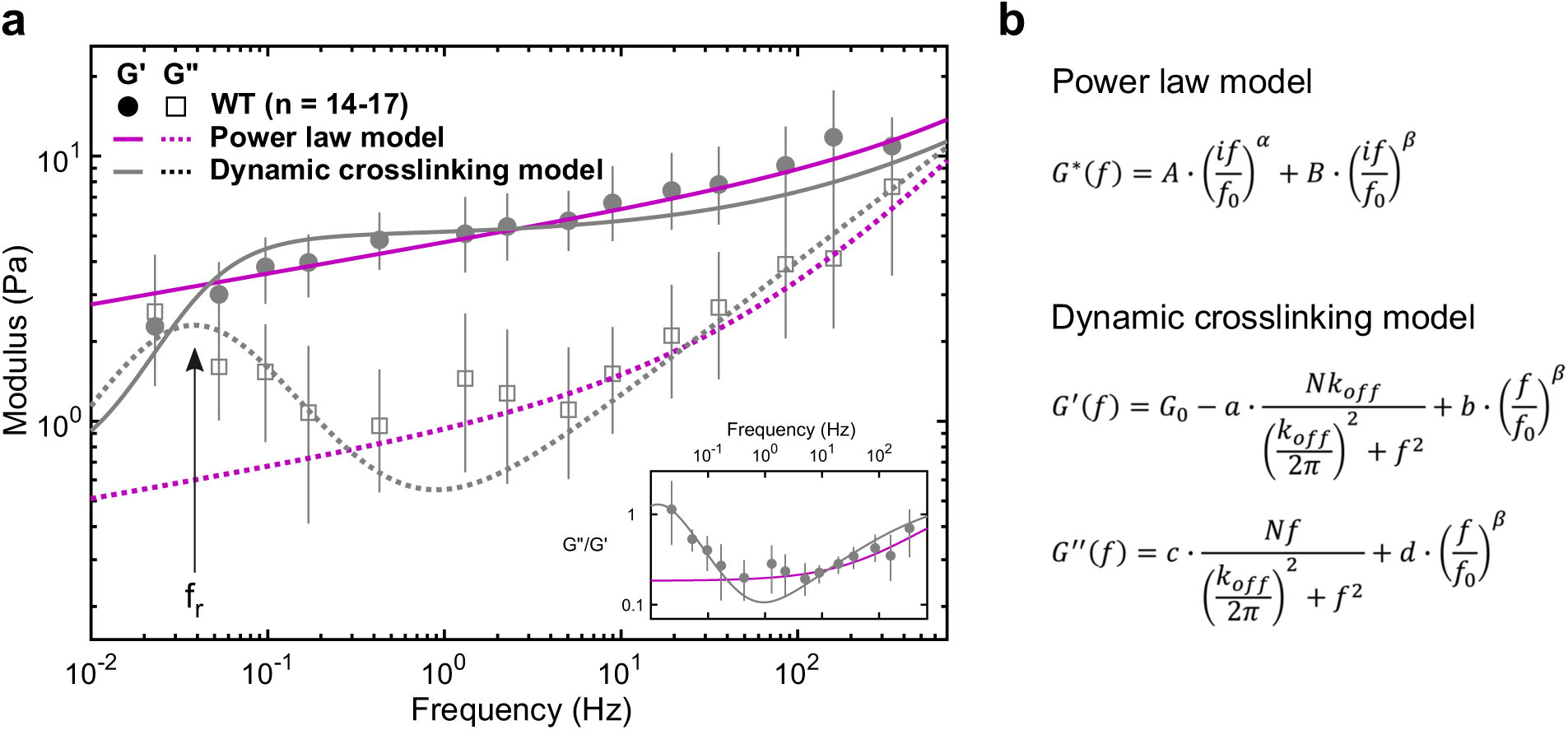
The viscoelastic properties of the cytoplasm show relaxation dynamics at lower frequencies. **a**, Elastic (G’) and viscous (G’’) moduli of WT fibroblasts (mean and 95% confidence interval from 1,000 bootstrap samples, see Methods). The fluid-like transition (i.e. relaxation) is visible at frequencies below ∼1 Hz: reading from right to left, there is a local G’’ minimum followed by a local G’’ maximum along with a drop in G’, also evident in the loss tangent (inset), with a minimum at ∼1 Hz and a steady rise with decreasing frequency. The dynamic crosslinking model (gray) captures the dynamics at lower frequencies better than the power law model (magenta, see Supplementary Fig. 5), with the local G’’ maximum corresponding to the relaxation frequency, f_r_ = k_off_/2π (arrow). **b**, Analytical expressions of the models. From the power law model fit, α = 0.116 ± 0.021 (mean and standard deviation of the means) and β = 0.717 ± 0.062. From the dynamic crosslinking model fit, k_off_ = 0.24 Hz (see Fig. 4b, gray) and β = 0.509 ± 0.027.

The first term of the double power law model describes a soft glassy material (SGM) at longer timescales and the second term describes frequency scaling from entropic filament fluctuations at shorter timescales^13^ (Fig. 3b, top). The rheology of a SGM has been empirically observed as elastically dominant and weakly scaling with frequency with a constant loss tangent of ∼ 0.1^47^. In contrast, our data indicate the loss tangent of the cytoplasm varies at frequencies below ∼1 Hz and reaches 1.0, indicating an equal viscous and elastic contribution at low frequencies (Fig. 3a, inset). Deviations from the SGM model at long timescales were previously observed using magnetic twisting cytometry of beads attached to the cell cortex^32^, suggesting that despite the differences in mechanical properties between the cell cortex and intracellular environment, a different rheological model must be considered at these long timescales. A model that assumes mechanics are governed by the unbinding kinetics of a dominant crosslinker^28^ captures the data well (Fig. 3a, gray). The first term of the dynamic crosslinking model is derived from the Fourier transform of the exponentially decaying lifetime of crosslinked points. This first term decreases the plateau elastic modulus G_0_ and increases the amount of viscous dissipation in isotropically crosslinked actin networks^28^. The second term describes filament bending fluctuations at shorter timescales as before. The dominant crosslinker’s characteristic timescale, k_off_, appears as a local maximum in the viscous modulus (Fig. 3a) at the frequency of relaxation (f_r_ = k_off_/2π). Fitting this model to the measured moduli yields an estimate of k_off_ = 0.24/s (Fig. 4b) for WT fibroblasts. This estimate is in agreement with unbinding rates measured for several passive actin crosslinkers including alpha-actinin, filamen, and fascin^48,49^. The dynamic crosslinking model was derived from bulk rheology of actin polymer gels where the unbinding of crosslinkers was related to bulk properties through the assumption that G_0_ ∼ N^x^ with x = 1 (where N is the number of crosslinked points) in a homogeneous and isotropically crosslinked network^28^. However, the cytoplasm is likely a composite network with the coexistence of bundled, branched, and isotropically crosslinked actin filaments, despite our efforts to probe the most homogenous region of the cytoplasm (Fig. 1c,d). Additionally, cell orientation and deformation axis also result in anisotropic mechanical properties^9^. The same model with G_0_ ∼ N^x^ and x not equal to 1 may account for some of this anisotropy. In addition, in bulk rheology, the shear stresses are assumed to be evenly distributed at the microscale. Thus, the resulting mechanical response is a summation of the contributions of individual crosslinkers across the sample. However, in microrheology, an increased viscous dissipation from the depletion of crosslinked points in the vicinity of the probe is expected, causing a crossover of elastic and viscous moduli at lower frequencies^28^. Thus, it may be possible to further improve the fit to the model by including a stronger frequency-dependent loss of crosslinking points.

**Figure 4.**
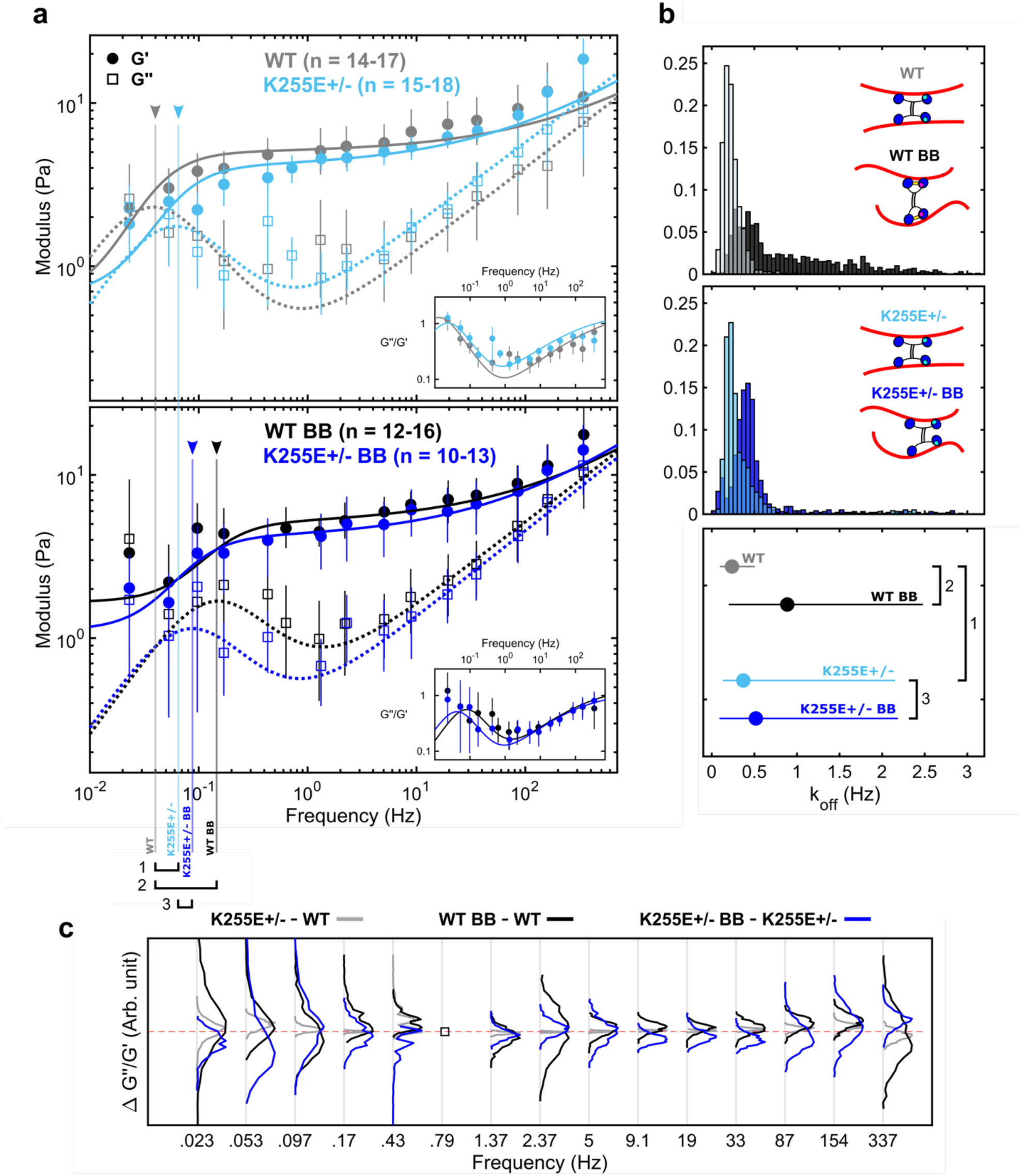
Relaxation dynamics after alteration of a-actinin 4 binding kinetics and myosin II disruption. **a**, Elastic (G’) and viscous (G’’) moduli of WT cells and ACTN4 K255E+/- mutant cells under control (top) and 50µM Blebbistatin (BB) -treated (bottom) conditions (mean and 95% confidence intervals from 1,000 bootstrap samples) along with the best fit to the dynamic crosslinking model. Untreated WT cells (gray) and K255E+/- cells (light blue) show similar dynamics, with little shift (1) in f_r_ (arrow head and vertical line). The f_r_ shift to the right from WT to WT BB (black) is apparent (2), indicating faster relaxation. The f_r_ shift from K255E+/- to K255E+/- BB (blue) is less apparent (3). **b**, Unbinding rate (k_off_) distributions (top and mid) and mean with 95% confidence interval (bottom) obtained from the bootstrap fits. The untreated WT and BB-treated WT cells show different k_off_ distributions, consistent with WT ACTN4’s binding kinetics. ACTN4 has two principal actin binding sites (inset schematic, blue heads) and a third putative actin binding site (cyan) that is exposed upon tension. Under reduced tension, the WT ACTN4 is in its closed conformation due to the presence of a crossbridge (yellow). The untreated K255E+/- and BB-treated K255E+/- cells show similar k_off_ distributions, consistent with the tension-independent binding kinetics. k_off WT_ = 0.24 Hz, k_off WT BB_ = 0.89 Hz, k_off K255E_ = 0.37 Hz and k_off K255E BB_ = 0.52 Hz (mean and 95% confidence interval) are obtained (bottom). The difference between these unbinding rates are proportional to the differences in f_r_ numbered in **a. c**, The difference between loss tangents (ΔG’’/G’) indicates that the mechanics of WT cells are tension sensitive while the K255E mutation reduces sensitivity to tension. The empty square at 0.79 Hz indicates a frequency point that is only present for one of the two conditions, resulting in no quantifiable difference. Distributions shifted above zero (red dashed line) indicate an increase in the loss tangent. ΔG’’/G’ between untreated K255E+/- and untreated WT cells (gray) show narrow distributions about zero, indicating little differences between the two conditions. ΔG’’/G’ between BB-treated WT and untreated WT cells (black) show a general upward shift, indicating that the BB-treated WT cells are more viscous compared to untreated WT cells. ΔG’’/G’ between BB-treated K255E+/- and untreated K255E+/- cells (blue) do not show a general trend, as indicated by the similar number of upward and downward shifts in distributions. At 0.43 Hz, the black curve is shifted upwards while the blue remains slightly below zero. This indicates an earlier transition to the fluid-like regime when WT cells are treated with BB compared to when K255E+/- cells are treated with BB. This upwards shift is consistent below 1 Hz, except at 0.097 Hz, indicating an overall increased fluid-like behavior at long timescales upon BB treatment in WT cells compared to K255E+/- cells.

Together, our data along with the fit to the model suggest that dynamic crosslinking of the cytoskeleton results in a fluid-like transition of the cytoplasm. By modulating the crosslinking dynamics of key crosslinkers, the cell could tune its viscoelastic properties in response to biochemical or mechanical cues. Correspondingly, dysregulation of cytoskeletal crosslinking may contribute to disease^50^. To investigate the role of actin crosslinker binding kinetics in governing cytoplasmic mechanics, we probe the viscoelastic properties of cells heterozygous for an ACTN4 mutation. ACTN4 is a ubiquitous crosslinker that dynamically crosslinks actin filaments with k_off_ ∼0.4 Hz^48^ and is involved in the formation of actin bundles^51^. The K255E mutation has been associated with focal segmental glomerulosclerosis (FSGS)^52^. In FSGS, podocytes are less able to resist cyclic loading, suggesting a dysregulation of the mechanical properties of the cytoskeleton^53^. Indeed, this mutation alone reduces cell motility while increasing cellular forces in heterozygous cell lines through an interplay with tension-generating myosin II motors^50^. Increased contractility is believed to be due to the exposure of a cryptic actin binding site in the K255E mutant, resulting in its increased affinity of for actin (i.e. slower k_off_)^51^. In WT ACTN4, the cryptic actin binding site is thought to only be exposed when the crosslinker is under tension, resulting in a catch-bond behavior^24^ (Fig. 4b, top inset). The difference in the unbinding rates of the WT ACTN4 and the K255E mutant results in a shift of f_r_ in purified networks^24,25,31^. In contrast, our results show no significant difference in the mechanical properties of WT and K255E+/- cells under normal conditions (Fig. 4a), possibly due to lower expression of K255E mutant and other mechanical anisotropies in the region probed. Overall, with k_off WT_ = 0.24 Hz and k_off K255E_ = 0.37 Hz (Fig. 4b) from the average best fit, our results suggest similar crosslinking dynamics in the WT and K255E+/- cells under normal conditions (Fig. 4b, top and bottom insets).

To investigate the role of tension on ACTN4 binding kinetics, we treat the cells with 50 µM Blebbistatin. Blebbistatin reversibly locks myosin II in the low-affinity ADP-P_i_ state, reducing tension in the actin cytoskeleton^54^. With 50 µM Blebbistatin, a sufficient number of cells (Supplementary Fig. 7) do not exhibit gross morphological changes and remain viable for mechanical measurements. Surprisingly, we do not observe significant softening of the cytoplasm (Fig. 4b) as reported previously^5^. This may be due to differences in Blebbistatin treatment response levels inducing more local heterogeneity, as depicted by the greater variability in the data (Fig. 4a, bottom) and the wider distribution of k_off_ (Fig. 4b, top), or differences in the *in situ* calibrated k_trap_ and β_pd_ values (Supplementary Fig. 2). It is also likely that the local cytoplasmic microenvironment is less sensitive to actomyosin activity, and thus Blebbistatin treatment, compared to on the cortex where a much more dramatic effect on magnitude is expected^14,55^. While the mechanics at fast timescales are not affected, at long timescales we observe a ∼3-fold faster transition to fluid-like behavior in Blebbistatin-treated WT cells, with k_off_ increasing from 0.24 Hz to 0.89 Hz in the presence of Blebbistatin (Fig. 4a, bottom). Interestingly, k_off_ in K255E+/- cells remained similar, with k_off K255E_ = 0.37 Hz and k_off K255E BB_ = 0.52 Hz. The results suggest that with reduced tension (through myosin II inhibition), actin binding kinetics differ in a manner that is consistent with the differences between the binding kinetics of WT and K255E ACTN4 (Fig. 4b, top and bottom insets). While tension levels vary between cells and within a single cell^56^, it is expected that, on average, the tension levels are reduced with Blebbistatin treatment. The active crosslinker myosin II may also directly play a role in regulating the observed mechanics as shown previously in purified networks^57,58^ at intermediate and long timescales. The effect of crosslinking dynamics on relaxation is also readily apparent in the differences in loss tangent (Fig. 4c). Latrunculin A (Lat A) depolymerizes actin filaments by sequestering free actin monomers^59^. Interestingly, treating WT cells with Lat A shifts the relaxation to faster timescales (Supplementary Fig. 6), suggesting actin filaments disruption plays a similar role in relaxation as the inhibition of intracellular tension^7^. In Lat A-treated cells, the presence of fewer, shorter actin filaments is expected to reduce the ability of ACTN4 and myosin II to interact with and crosslink actin filaments, reducing the tension generated and stored in the network^58^. This reduced number of crosslinked points does not alter the frequency of relaxation in purified networks^25,28,29^, suggesting that the shift in relaxation that we observe in Lat A-treated WT cells results from changes in binding kinetics, similar to what is observed with Blebbistatin treatment. In both Blebbistatin - and Lat A-treated cells (Fig. 5 and Supplementary Fig. 6), the loss tangent indicates that the network is more fluid-like at lower frequencies, consistent with previous observations using AFM^7^. However, similar to Blebbistatin treatment, and in contrast with previous studies^7,18^, we observe negligible differences in the magnitudes of G* with Lat A treatment^55^. Intermediate filaments, another component of the cytoskeletal network, contribute approximatively half of the mechanical resistance in the intracellular environment^10^ and thus help maintain mechanical integrity even when the actin cytoskeleton is disrupted. Furthermore, cells that show severe signs of actin cytoskeleton disruption are not viable for mechanical measurements. Consequently, healthier cells (Supplementary Fig. 7) were chosen preferentially to yield more consistent and quantifiable results, which may partly explain the similar magnitudes observed under Blebbistatin and Lat A treatment. Overall, our results show a shift in relaxation to faster timescales when myosin II is inhibited, particularly in the case of WT cells, suggesting an earlier transition to the fluid-like regime consistent with the tension-dependent differences in binding kinetics between WT and K255E ACTN4.

While perturbations to the crosslinked actin network modulate the transition to fluid-like behavior, the mechanics at timescales >1 Hz are largely unaffected. Intermediate filaments^10^ and microtubules^60,61^ also contribute to the mechanical response of the cytoplasm, particular in the cell interior^55,4^. Further studies are needed to dissect the relative roles of actin, microtubules, and intermediate filaments in governing cell mechanics.

Mechanical properties play an important role in the regulation of many cellular functions and allow the cell to be solid enough to provide structural support but also fluid enough to reorganize and perform dynamic functions. By developing a novel method to measure intracellular mechanics, our measurements reveal relaxation of the cytoskeleton at long timescales across all conditions, pointing to a fundamental mechanical property of the cytoplasm. In addition, a model that includes the binding kinetics of a dominant actin crosslinker captures these relaxation dynamics well. However, while our data suggest a relationship between mechanical relaxation and the crosslinking dynamics of ACTN4, many other crosslinkers and motor proteins contribute and it is unlikely that ACTN4 alone governs the viscoelasticity at these timescales. Other cytoskeletal components with their associated proteins may also contribute to cytoplasmic relaxation through mechanisms that are yet to be identified. Overall, we performed calibrated mechanical measurements in the cytoplasm of living cells using optical tweezers and related the relaxation dynamics observed to the binding kinetics of ACTN4. Future work focusing on other crosslinkers and other components of the cytoskeleton, as well as on simplified *in vitro* systems, will add to the results reported here and provide a more complete understanding of the mechanisms governing the fluid-like transition of the cytoplasm.

## Acknowledgments

The authors thank Gary Tom and Giancarlo Szymborski for their work on the control software for the optical trap, Chris Sitaras for helpful discussions and assistance in PEG-coating of the beads, and both Chris Sitaras and Ajinkya Ghagre for bulk rheology data. The authors also thank Malina K. Iwanski, Abdullah R. Chaudhary and Linda Balabanian for helpful discussions and valuable feedback on the manuscript. AGH acknowledges support from NSERC (RGPIN-2014-06380) and the Canadian Foundation for Innovation. AJE acknowledges support from NSERC (RGPIN/05843-2014, EQPEQ/472339-2015, RTI/00348-2018), CIHR (Grant # 143327), and the Canadian Foundation for Innovation (Project #32749).

## Author Contributions

A.G.H., A.J.E., H.K.H. and L.C. designed the research. L.C. and H.K.H. developed experimental protocols and analytical tools. L.C. and A.R.C. performed experiments. L.C. analyzed the data. A.J.E. provided cells and reagents. L.C. and A.G.H. wrote the paper.

## Additional information

Supplementary information

## Methods

### Cell culture and preparation

Two immortalized dermal fibroblasts cell lines are used^50^. Cells are cultured in DMEM media (Thermo Fisher Scientific), supplemented with 10% FBS (Thermo Fisher Scientific) and 1% Glutamax (Thermo Fisher Scientific). Cells are passaged at 80% confluency and seeded on glass coverslips 24h before the experiment at low enough confluency to minimize cell-cell contacts. 500 nm diameter fluorescent latex beads (Thermo Fisher Scientific) are PEG-coated following previously published protocols^62^. Prior to the experiment, the beads are washed and resuspended in hypertonic media (∼10^9^ beads/mL in 10% PEG 35000, 0.25 M sucrose in complete DMEM media). The cells are incubated for 1 hour with the hypertonic media to allow the beads to be phagocytosed. A gentle osmotic shock is administered to the cells to burst endogenous membranes formed around the internalized beads, minimizing further interactions of the bead with proteins of the endocytic pathway^33,64,65^. This shock is applied by incubating the cells with a hypotonic solution for 3 minutes. The cells are then left in the incubator in regular complete media for one hour to recover from the shock. Tracking the difference in mechanics over time of cells hypotonically shocked at the same time shows no significant changes (Supplementary Fig. 8), suggesting that the one-hour recovery period is enough. The coverslips are mounted on microscope slides using double-sided tape to form a flow chamber. In some experiments, media is replaced with complete media supplemented with 50 µM BB or 100 nM Lat A. Measurements start 15 minutes after treatment and cells are kept at 37 °C using a heater (World Precision Instruments) in a custom-built environmental chamber during all optical trapping experiments.

### Optical trap

The optical trap is installed on an inverted microscope (Eclipse Ti-E: Nikon, 1.49 NA oil-immersion 100x-objective). The near-IR 1064 nm laser beam (10 W, IPG Photonics) is expanded to overfill the back aperture of the objective. The bead position relative to the laser center is measured using a lateral effect photodiode (Thorlabs) positioned conjugate to the objective’s back focal plane. The laser is steered using 2-axis AODs (AA Optoelectronic) controlled through a field-programmable gate array and custom LabVIEW programs (National Instruments). A multifrequency (∼18 frequencies) excitation input wave is applied to a single intracellularly located bead per cell. The frequencies range from 0.02 Hz to 1300 Hz (moduli obtained from 0.02 to 500 Hz), with corresponding amplitudes ranging from ∼50 nm to ∼1 nm. The frequencies of oscillation are selected to avoid harmonics. The amplitudes are empirically chosen to yield a linear response from the cytoplasm. No significant harmonic generation was observed from the amplitudes applied, indicating that the measurements were performed in the linear regime^63^. Measurement are performed on beads located in the most mechanically homogenous region of the cell, midway between the cell periphery and perinuclear region where distinct actin cytoskeleton organization exists.

### Optical trap data analysis

Data is acquired at 200 kHz downsampled to 20 kHz or directly acquired at 20 kHz (Supplementary Fig. 1). k_trap_ and β_pd_ are obtained for each individual measurement as previously described^35^. In brief, the frequency response function (FRF) is obtained from the cross spectral density of the bead displacement and AOD using Welch’s method in MATLAB. The window length is chosen at each individual frequency to be an integer multiple of the period of the input frequency to avoid leakage^34^. The integer multiple, is chosen to obtain the best trade-off between frequency resolution and averaging. A simplified viscoelastic model is simultaneously fit to the FRF and the power spectrum of the thermal response to obtain k_trap_ and β_pd_^35^. The elastic and viscous moduli are directly obtained from the FRF knowing k_trap_ and β_pd_^38^. See Fig. 2 and Supplementary Information for more details.

### Statistics and fits

1000 bootstrap samples are drawn from the data and each bootstrap sample is fit to either the dynamic crosslinking model or the power law model, allowing distributions of fit parameters to be obtained for each condition. The power law model consists of two frequency terms raised to the power of α and β respectively (see Fig. 2b), added together and scaled with constants A, B and f0. The real and imaginary part yield the elastic and viscous modulus respectively. The dynamic crosslinking model includes a k_off_-dependent term. G_0_ is the plateau elastic modulus, N is the number of crosslinked points, a, b, c, d and f_0_ are scaling constants. The fits are done using the nonlinear least squares MATLAB function. Due to differences in excitation inputs applied across experiments, the number of experiments (n) used to compute the mean bootstrap moduli at each frequency varies slightly. To compare the two model fits, Bayesian Information Criteria (BIC) are used. Fitting bootstrap samples allows statistics to be directly obtained and also allows a more representative comparison of the fit parameters between conditions, as oppose to directly fitting the mean of the data. See Supplementary Information for more details.

## Supplementary Figures

**Supplementary Figure 1.**
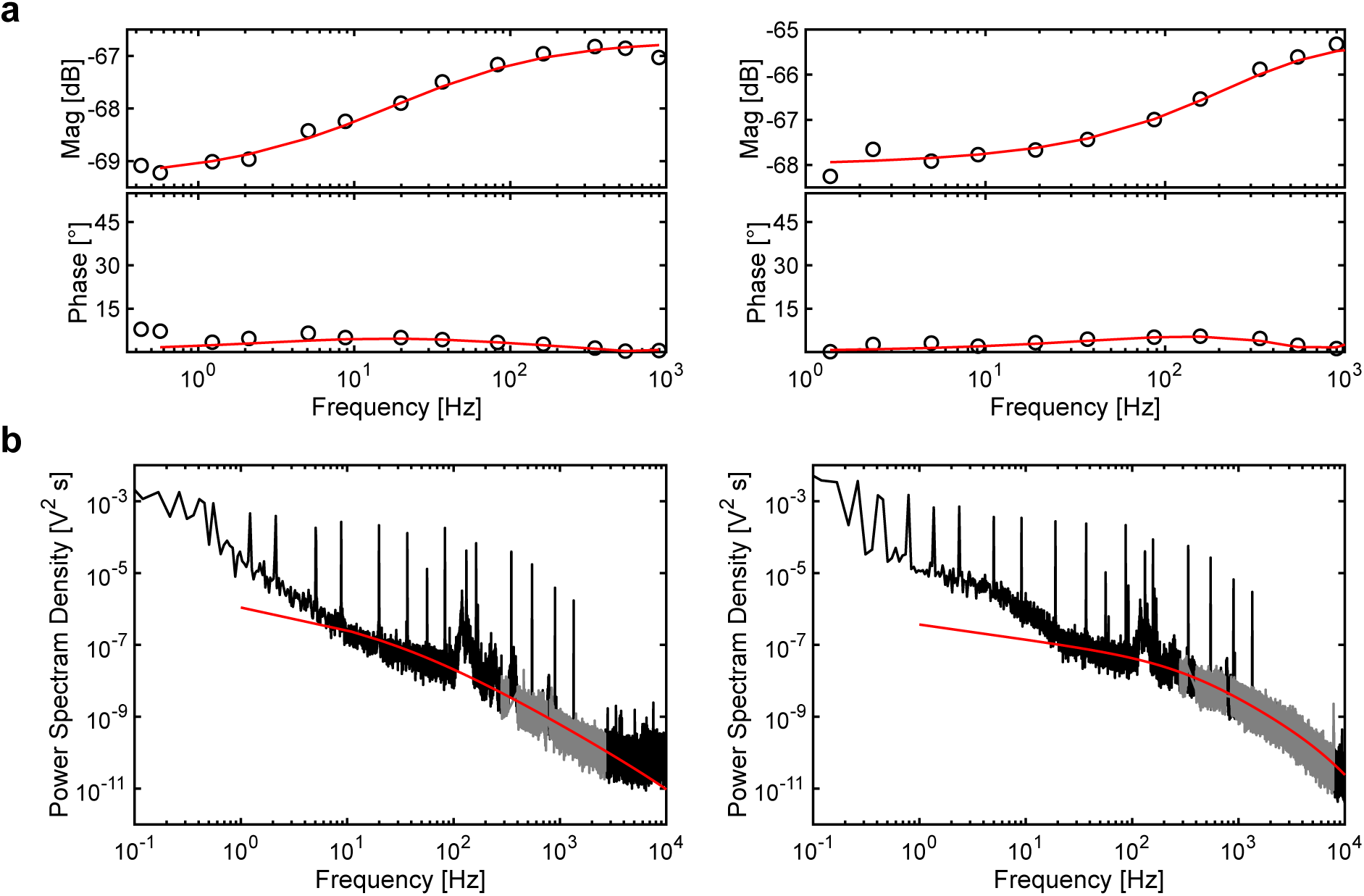
Additional trap calibration examples. **a**, Frequency response function (FRF) from two representative samples. The best fit to the FRF of the simple viscoelastic model described above is shown in red. **b**, Power spectrum from the corresponding sample as in **a**. The best fit to the power spectrum of the model is shown in red. On the left, direct data acquisition at 20kHz is used, aliasing can be seen at frequencies above ∼2500 Hz with the noise floor being reached at frequencies past ∼3000 Hz. The part of the power spectrum that is fit is shown in grey and stops before reaching the aforementioned frequencies. On the right, data acquisition at 200 kHz downsampled to 20 kHz through averaging is used. A filtering effect from the averaging can be seen starting at ∼2500 kHz with a second apparent corner frequency. This filtering was corrected during the fit by adding a low pass filter with corresponding corner frequency.

**Supplementary Figure 2.**
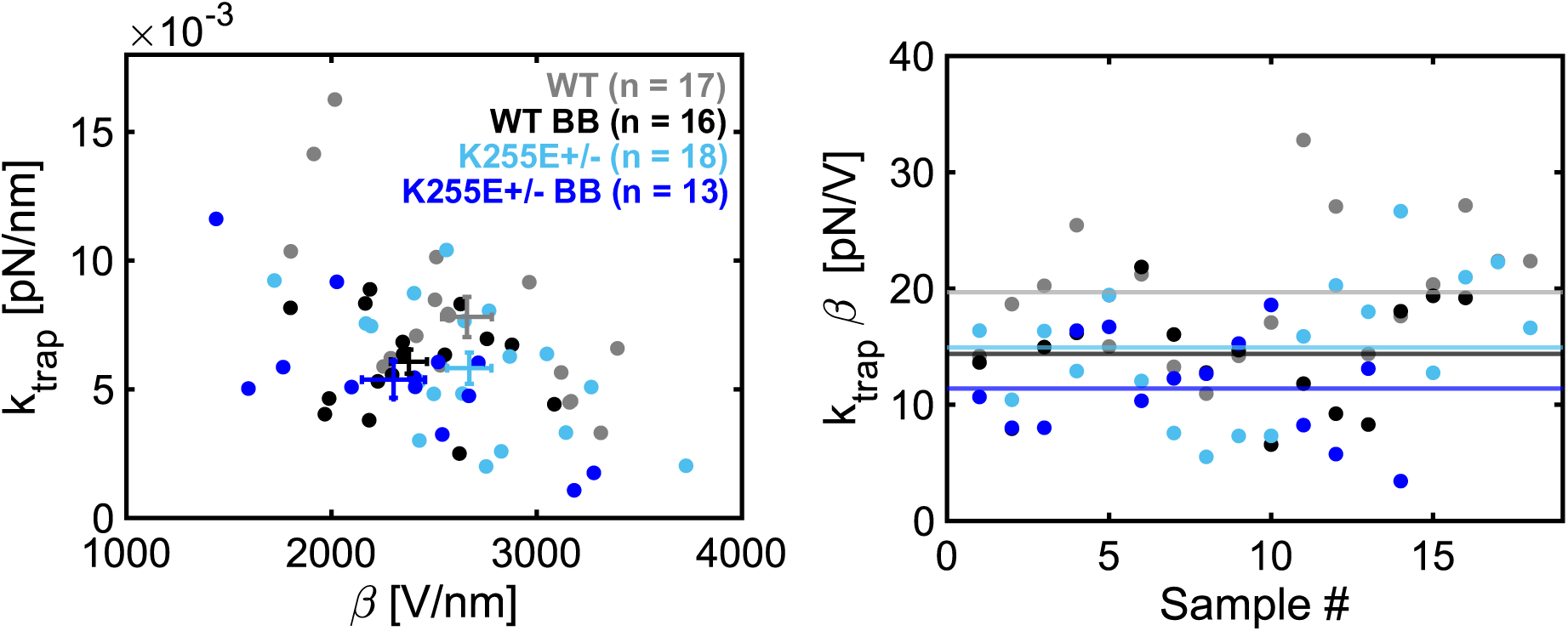
Calibrated k_trap_ and β_pd_ for all experiments. **Left:** k_trap_ and β_pd_ vary between cells and experimental conditions. Error bars indicate SEM. **Right**: the product of k_trap_ and β_pd_ clusters around 15 pN/V. The mean values are plotted (lines) and indicate that most of the k_trap_ x β_pd_ fall between 10 and 20 pN/V.

**Supplementary Figure 3.**
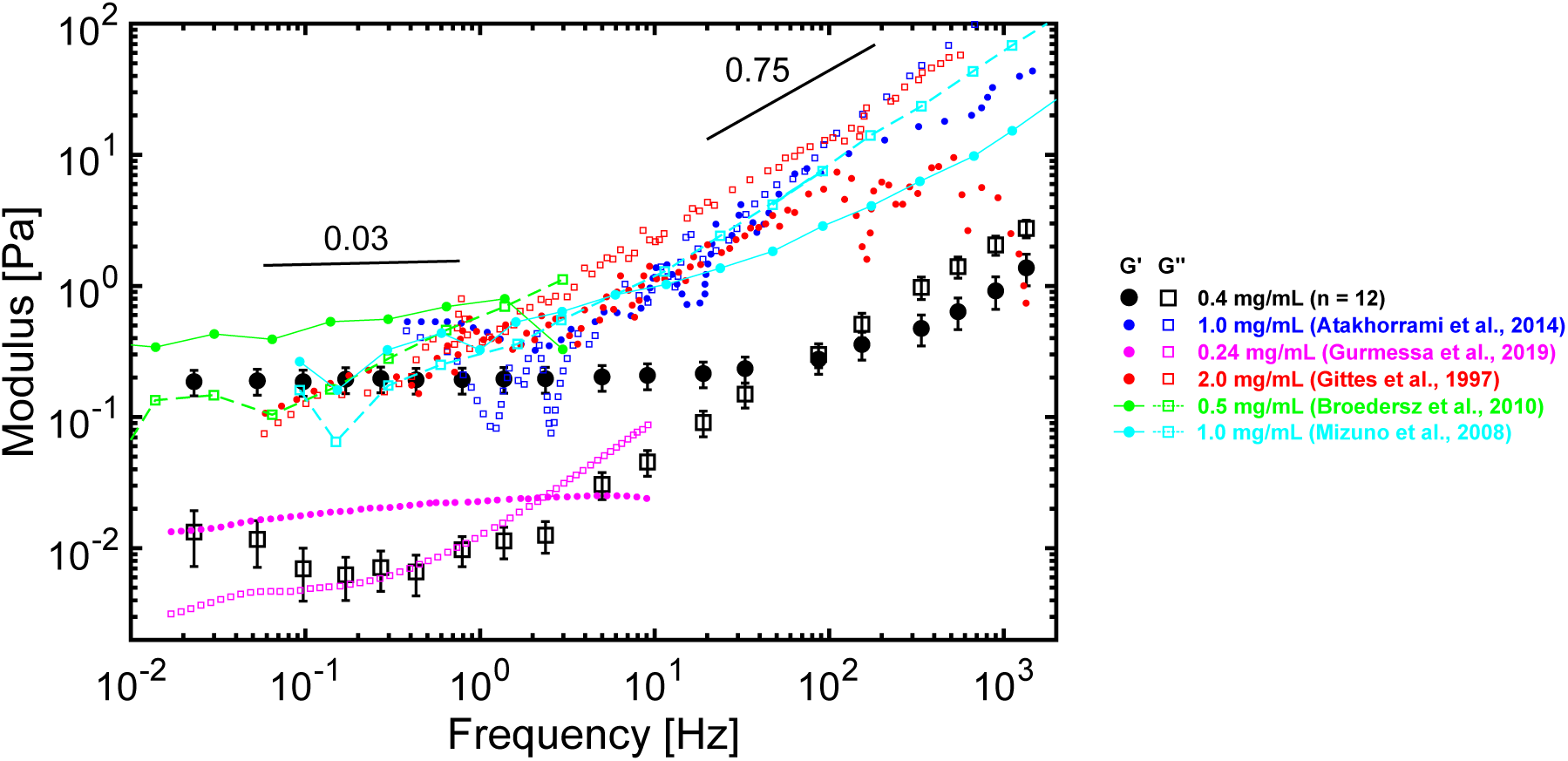
Viscoelastic response of purified actin networks. The elastic (G’, black) and viscous (G’’, black) moduli obtained from microrheological measurement with the optical trap and *in situ* calibration in 10 µM (∼0.4 mg/mL) actin networks. Below ∼30 Hz, the elastic modulus weakly scales with frequency (∼ f^0.03^) and above ∼50 Hz, both G’ and G’’ strongly scale with frequency (∼ f^0.75^). At frequencies below 1 Hz, G’ clearly dominates, indicating a mostly elastic response and no apparent viscous relaxation. These results are in excellent agreement with previous reports on actin networks mechanics at similar actin concentrations (blue^1^, purple^2^, red^3^, connected green dots^4^ and connected cyan dots^5^, both in terms of frequency scaling and magnitudes.

**Supplementary Figure 4.**
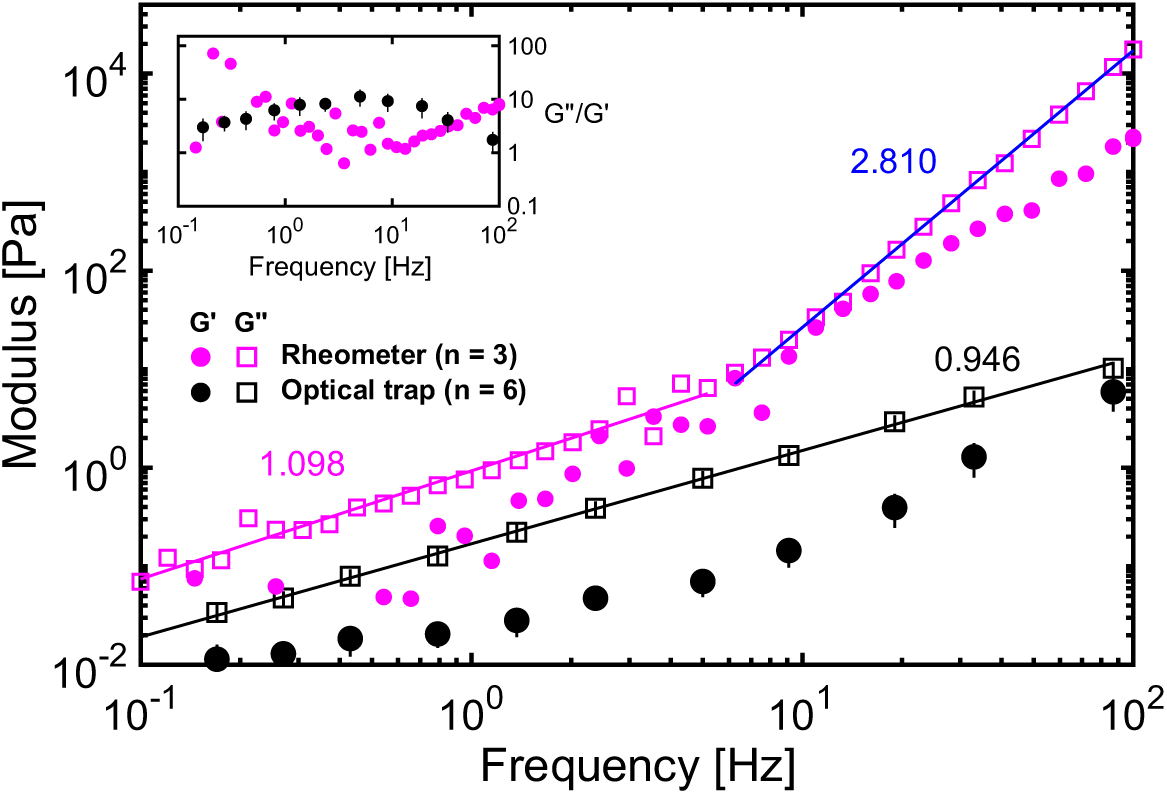
Micro and bulk rheology of 20% Polyethylene Glycol (PEG). The elastic (G’, solid circles) and viscous (G’’, open squares) moduli from a rheometer (magenta, MCR 302, Anton Paar) are compared to moduli obtained from the optical trap (black). The values obtained from bulk rheology show G’’∼ f^1.098^ (magenta line, magenta number) at frequencies below ∼7 Hz, in agreement with the G’’ scaling measured from microrheology, with G’’∼ f^0.946^ (black line, black number). At frequencies past ∼7 Hz, a stronger G’’ scaling is observed for the bulk data, with G’ ∼ f^2.810^ (blue line, blue number). This is likely due to inertial effects coming from the instrument and the sample itself that can show an apparent increased frequency dependence at higher frequencies^6^. For optical trapping measurements at the microscale, inertial forces remain negligible at these timescales. In terms of magnitude, the data from the rheometer show significantly greater viscous and elastic moduli across the whole frequency spectrum. This is likely due to differences between the length scales investigated in bulk and microrheology that result in a different mechanical response to deformation^7,8^. From the loss tangent (G’’/G’, inset), before 1 Hz, the rheometer is unable to probe the elastic modulus reliably, resulting in very low elastic moduli (out of the plotted region in the main plot) and jumps in the loss tangent. Between 1 Hz and 10 Hz, the rheometer data show lower loss tangent, indicating that there is a greater elastic contribution, likely from surface tension^6^. Above 10 Hz the loss tangent increases due to inertial effects as discussed above.

**Supplementary Figure 5.**
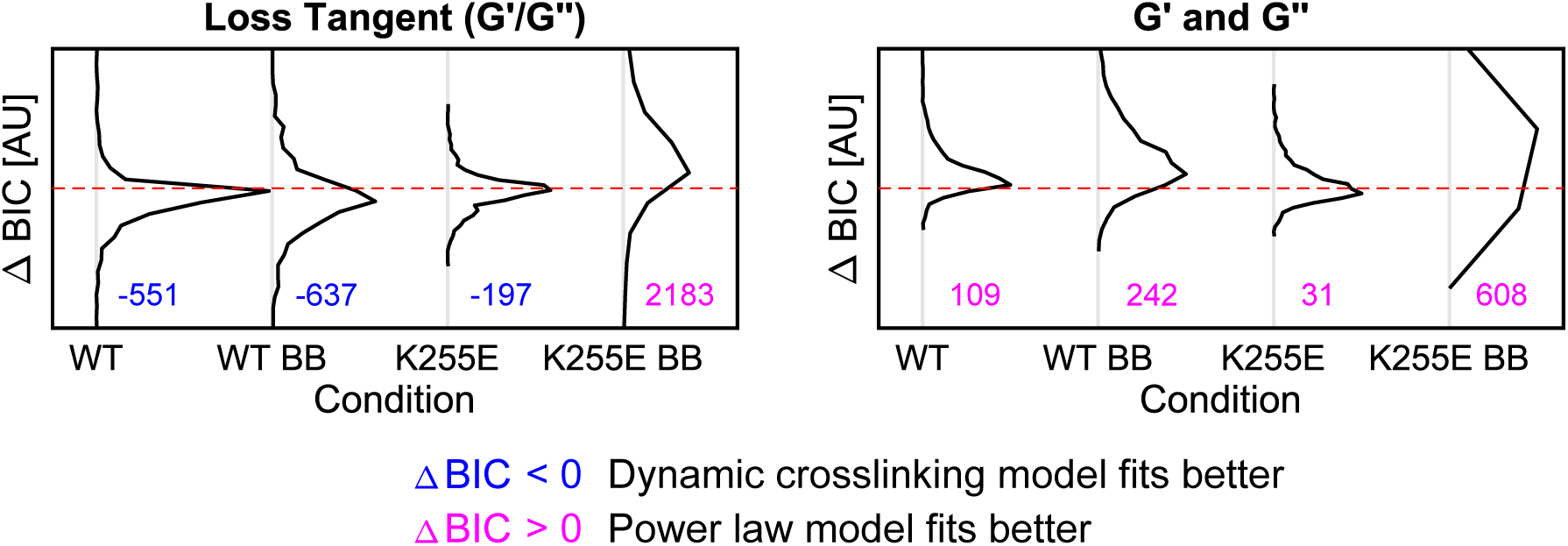
Difference in Bayesian Information Criteria (BIC) values. **(A)** The distribution of the difference in BIC (ΔBIC) values from the loss tangent (G’’/G’) fit between the dynamic crosslinking model and power law model (see Fig. 3b) is obtained at each bootstrap sample for all four conditions and plotted vertically (y-axis and x-axis are both arbitrary units). The number next to the distribution indicates the mean ΔBIC values. A downward shift from zero (dashed red line) indicates a smaller BIC value for the dynamic crosslinking model, i.e. a better fit using that model. Wild type (WT), Blebbistatin(BB)-treated WT (WT BB) and mutant (K255E+/-) cells show a downward shift suggesting that the dynamic crosslinking model fits better (blue). Surprisingly, K255E+/- BB show an upward shift, suggesting that the power law model captures the loss tangent better (purple), possibility due to the inability of the dynamic crosslinking model to capture the slowest and fastest frequencies. **(B)** The distribution of ΔBIC values obtained from the fit to G’ and G’’ show an upward shift for all conditions, suggesting that the power law model captures G’ and G’’ better (purple). This may be due to single frequency point magnitude shifts that contributes to increased error terms. The loss tangent is less sensitive to these sources of error and is likely better suited for BIC values comparison. Overall, while the ΔBIC values distributions comparing the power law and dynamic crosslinking model are not statistically significant, the power law model is unable to capture the relaxation dynamics at low frequencies as seen in the loss tangent plots (see Figs. 3a and 4a in main text and Supplementary Figure 6) where a distinct V-shape is observed.

**Supplementary Figure 6.**
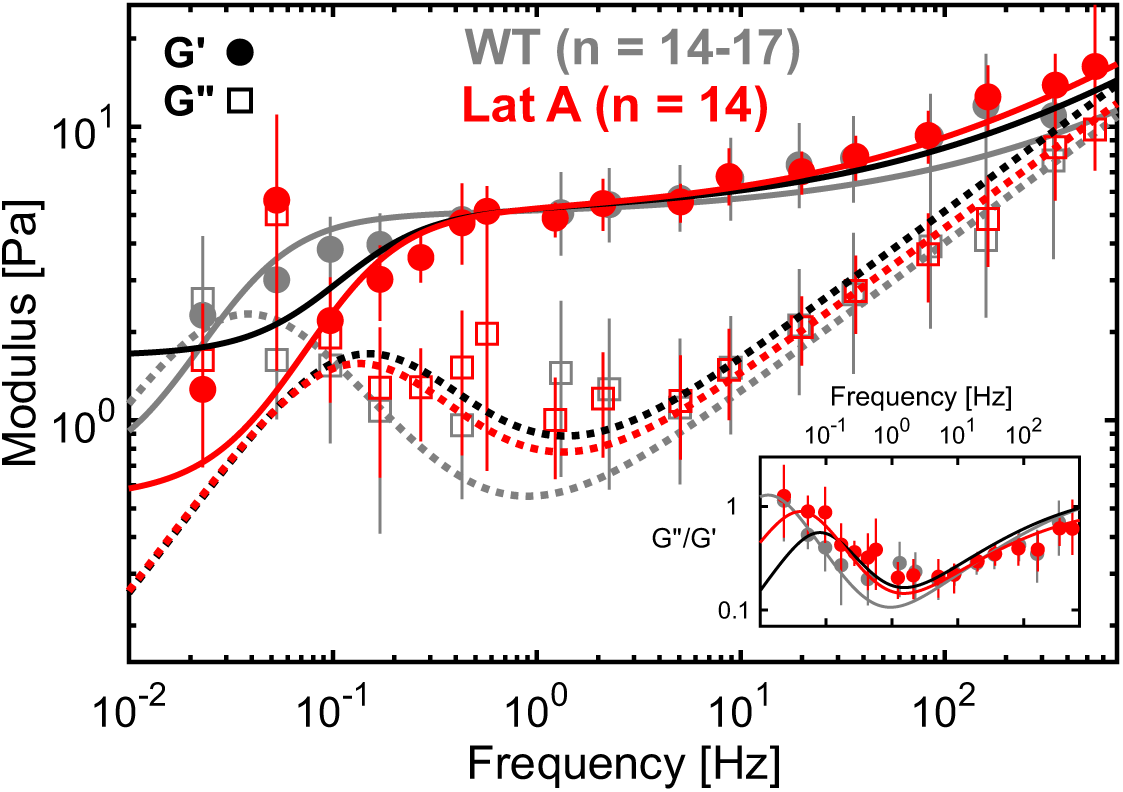
Relaxation dynamics after actin filament disruption. The elastic (G’, solid circles) and viscous (G’’, open squares) moduli from untreated WT (gray) and cells treated with 100nM Latrunculin A (Lat A, red). The best fit to the Blebbistatin-treated WT cells is shown in black for comparison. The Lat A-treated cells show faster relaxation (k_off_ = 0.80 Hz) compared to untreated WT (k_off_ = 0.24 Hz), as indicated by the earlier drop in G’ and earlier rise in G’’ at ∼0.4 Hz. This is also visible in the loss tangent (inset), with a minimum observed at ∼1 Hz, and an earlier rise in loss tangent with decreasing frequency, overall indicating an earlier transition to the fluid-like regime. Lat A-treated cells also show very similar relaxation compared to the Blebbistatin-treated WT cells (k_off_ = 0.89 Hz), as indicated by the near overlap between the red and black curves. Lat A concentrations greater than 100nM lead to complete cell death (data not show) while at 100 nM, only a small fraction of the cells showed severe signs of actin depolymerization and were avoided for mechanical measurements.

**Supplementary Figure 7.**
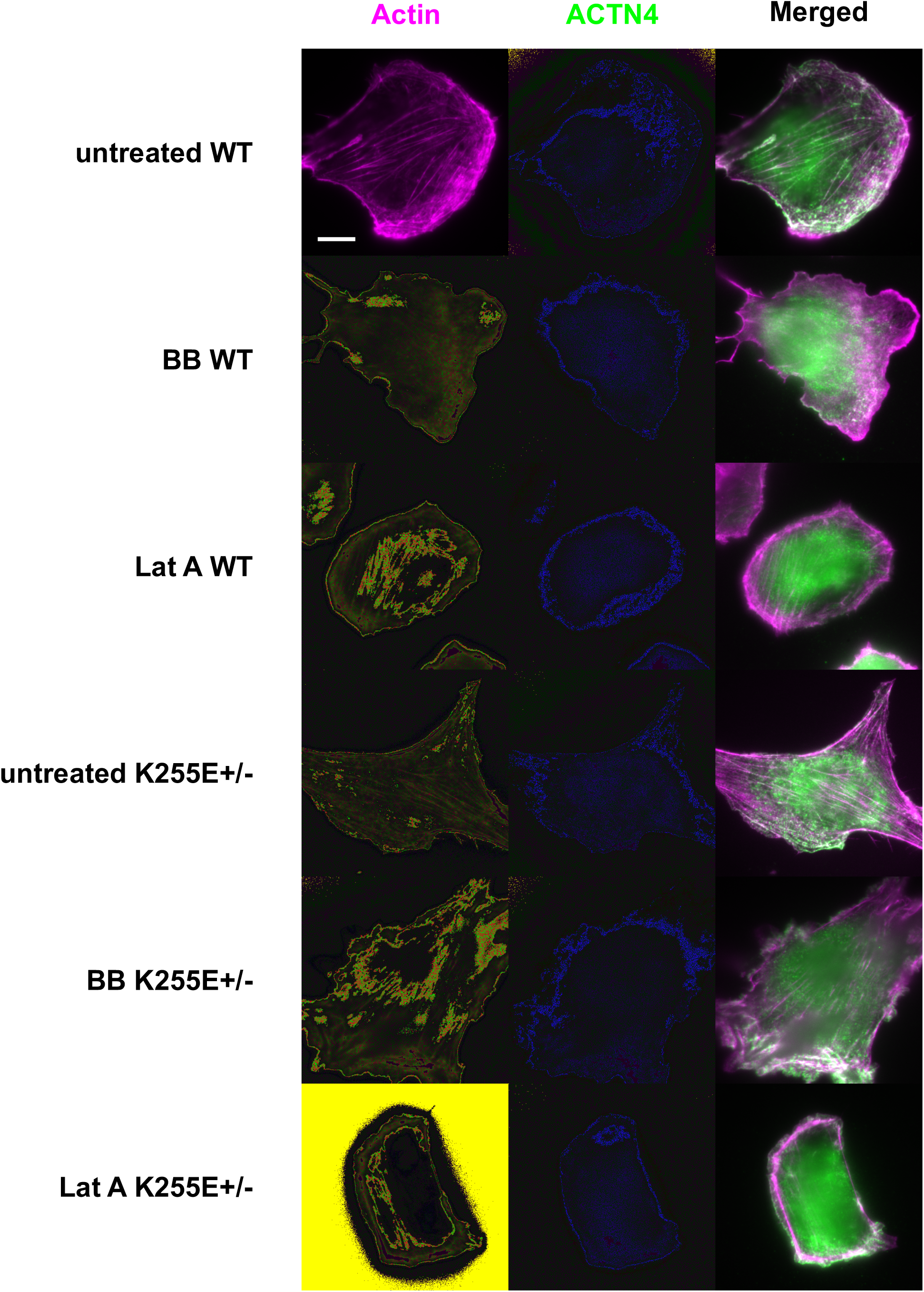
Immunofluorescent images of the WT and K255E+/- cells in various conditions. WT (top three rows) and K255E+/- mutant (bottom three rows) immunofluorescent images under normal conditions, after 50µM Blebbistatin (BB) treatment and after 100nM Latrunculin A (Lat A) treatment. ACTN4 was stained with 1:350 primary anti-ACTN4 (ab59468, abcam), incubated overnight at 4°C, followed by 1:350 secondary anti-rabbit (Anti-Rabbit Atto 488, Sigma-Aldrich) incubated for 30 mins along with 1:100 phalloidin (Alexa Fluor™ 647 Phalloidin, Thermo Fisher Scientific). Images show the maximum intensity projection of 3 Z stacks 200 µm apart acquired using near-Total Internal Reflection Fluorescence (near-TIRF) illumination. In normal conditions, K255E+/- cells are slightly larger and show more developed actin stress fibers compared to WT cells (untreated, actin column) in agreement with observations on homozygous K255E+/- mutant cells^9^. K255E+/- cells show less specific ACTN4 localization to both actin stress fibers and regions of increased actin enrichment (untreated, ACTN4 and merged column), suggesting that K255E ACTN4 does not preferentially bind to actin filaments under tension or active reorganization, in agreement with its binding kinetics. K255E+/- cells also show little preferential actin enrichment at the cell leading edge suggesting an impaired cell motility machinery, consistent with the mutant’s reduced motility reported previously^10^. For both WT and K255E+/- cells, Blebbistatin treatment reduces the number of visible stress fibers and causes highly irregular cell membrane shape at the cell periphery (Blebbistatin treatment, actin column), indicating a disruption in the actomyosin machinery. Under Lat A treatment, actin is most visible at the cell periphery (Lat A treatment, actin column), indicating that thick bundles and dense actin networks at the cell periphery are able to resist Lat A better compared to single or weakly bundled actin filaments in the cytosol, consistent with Lat A’s expected effect. Overall, we observe only subtle differences between the WT and K255E+/- mutant cells, which may also explain the subtle differences we measure in cytoplasmic viscoelasticity. Careful quantification of these actin and ACTN4 localization differences in the future would help in further interpreting the mechanical measurements. Finally, it is important to note that cells under both Blebbistatin and Lat A treatment only show mild signs of actin cytoskeleton disruption. Indeed, the concentrations of Blebbistatin and Lat A were chosen to cause a mild disruption of the actin cytoskeleton to ensure cells were viable for mechanical measurements and no large-scale morphological changes were observed.

**Supplementary Figure 8.**
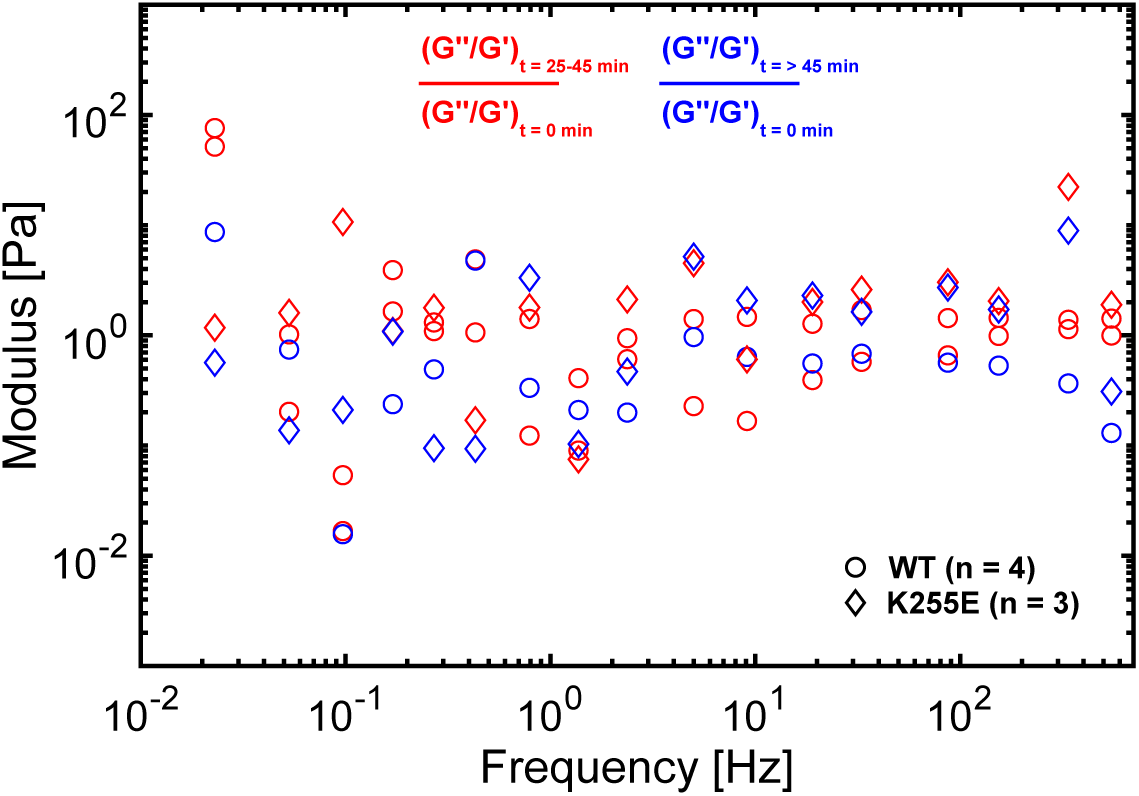
Loss tangent change over time of different cells after the one-hour recovery from hypotonic shock. Each point represents the change in loss tangent after an additional 25-45 mins (red, t = 25-45 min) or 45 mins (blue, t = > 45 min) has elapsed since the completion of the one-hour recovery (not shown for clarity, t = 0 min). Each point is the ratio of the loss tangent over its corresponding initial value as indicated at the top of the plot. A set of experiment with WT cells (circles) shows the change in loss tangent of two measurements performed after 25-45 mins (red circles) and one measurement performed after over 45 mins (blue circle). Within this set of experiments, the red and blue circles do not show any consistent patterns with the change in time. Similarly, within a set of experiment with K255E cells (diamonds) the red and blue diamonds also do not show any consistent patterns. Moreover, the changes in loss tangent of both experimental sets taken together (all red and all blue symbols) do not show any observable patterns and they cluster at ∼1, indicating little changes in mechanics over time. Both sets of experiments were performed under control conditions (no drug treatments).

## Supplementary Information

### Data handling

To collect data over several periods of excitation, a single measurement typically lasts 220 seconds or longer and active oscillations across all ∼17 frequencies are continuously being applied. The frequency response function (FRF), i.e. the transfer function H(ω), is obtained using Welch’s method, implemented in MATLAB with the “tfestimate” function. The Hamming window function is used and gives a slightly better response in terms of magnitude and coherence compared to the Hann window. The transfer function estimate is obtained at each input frequency separately with separate window lengths (NFFT). The window length is chosen to be an integer multiple of the input frequency to minimize signal leakage into adjacent frequency bins. The per cent overlap between windows is chosen based on empirical observations to yield the maximum magnitude and maximum coherence. At the lowest frequencies, less periods of excitation are contained in the measurement total time and the per cent overlap is thus chosen to be larger to allow more averaging over the ∼4 windows. At higher frequencies, the overlap is chosen to be 50% over the 6-8 windows. All windows and overlap amounts are chosen such that at most, ∼15 % of the data is truncated at the end. Indeed, as we have a finite sample length, truncation of the data occurs when the windows don’t exactly add up to the total length. Overall, the aforementioned window lengths and overlaps yield the best balance between frequency resolution and averaging. However, a more detailed and quantitative characterization may help for future application of this method. The magnitude and phase obtained are fit to the active part of the viscoelastic model discussed further below.

The power spectrum used for the thermal response is obtained using the same data. The peaks that correspond to the active oscillations are cropped out based on the known excitation frequencies. The remaining power spectrum represents the thermal response. For data that is acquired at 200kHz downsampled to 20kHz using a Field Programmable Gate Array, a pole is added to the viscoelastic model (corner frequency ∼ 3000 Hz) to account for this low pass filter coming from the downsampling by averaging. Sampling at 200kHz downsampled to 40kHz shifts this filtering to higher frequencies, confirming that the averaging is acting like a low pass filter (data not shown). Data acquired directly at 20kHz show a reduced signal to noise ratio compared to oversampled and averaged data, where the noise floor is visible on the power spectrum at high frequencies (Supplementary Fig. 1b). In these cases, the power spectrum is fit only up to the point where this upwards shift is observed.

### *In situ* calibration

From the mechanical circuit (Fig. 2b), for the case where the stage is stationary and by making the following substitution: Δx(t) = x_bead_(t)-x_trap_(t) and k_cyt_(t) = k_cyt,0_ + k_cyt,1_t^-(α-1)^, the following equation of motion can be obtained

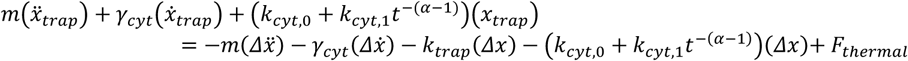

Substituting the cytoplasmic stiffness (k_cyt_) by the sum of the constant stiffness (k_cyt,0_) and the frequency dependent stiffness (k_cyt,1_t^-(α-1)^) observed in semiflexible polymer network allows the model to account for some of the viscoelastic properties of the cytoplasm11. Looking at the active part only (i.e. F_thermal_ = 0), neglecting inertial forces 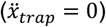 and taking the Laplace transform at steady state (s = jω), the following transfer function H(ω) = ΔX(ω) / X_trap_(ω) is obtained:

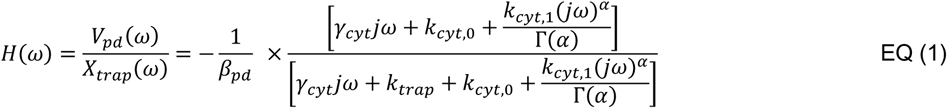

Where V_pd_(ω)β_pd_ = ΔX(ω) is used and represents the voltage from the photodiode and the photodiode calibration constant respectively. For visualization, if one ignores the complex cytoplasmic stiffness (k_cyt,1_ = 0), the numerator is a single zero proportional to 1/k_cyt,0_ and the denominator is a single pole proportional to the inverse of the total stiffness, 1/(k_trap_ + k_cyt,0_). The zero and pole are responsible for the two inflection points in the magnitude response, leading to the sigmoidal curve observed in the magnitude response obtained experimentally (Fig. 2b and Supplementary Fig. 1a). The addition of the frequency dependent stiffness shifts the zero and pole positions and changes the sharpness of the transition between the two^11^. The magnitude and phase of EQ (1) is what we fit to our experimental H(ω) obtained through *tfestimate* as outlined above.

Looking at the passive part only (X_trap_ = 0) and taking the Laplace transform at steady state as before, the following expression is obtained:

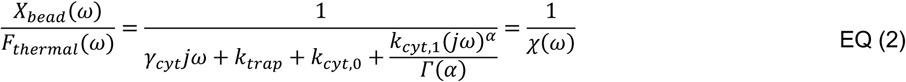

This complex response function *χ*(ω) can be related to the complex shear modulus through Stokes relation^3^

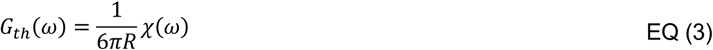

Here, G_th_(ω) represents the theoretical complex shear modulus based on the simple viscoelastic model, and R is the radius of the spherical probe. From the fluctuation-dissipation theorem, the power spectral density 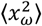 of the bead can be related to this response function *χ*(ω) or equivalently, to G_th_(ω) as follows^12^:

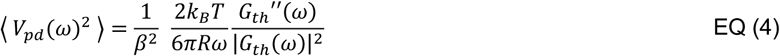

Where the following substitution has been done 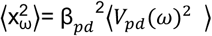, V_pd_ is the voltage measured by the photodiode, k_B_ the Boltzmann constant and T the temperature. EQ (4) is the equation fit to the power spectrum, with G_th_(ω) defined by EQ (2) through EQ (3). The corner frequency in the power spectrum is the result from the sum of the total stiffness of the system and corresponds to the pole in the magnitude response described above. α dictates the slope after the corner frequency, with α=1 corresponding to purely diffusive behavior and a slope of 2 in the power spectrum, and α<1 corresponding to constrained diffusion with a slope < 2 in the power spectrum. For more details on the different parameters of the model, see reference^13^.

Once the calibration is done, the complex shear modulus G(ω) of the real cytoplasm can be directly obtained using the experimental transfer function H(ω) of the measurement following reference^5^:

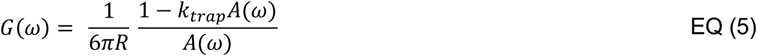

Where A(ω) is the apparent complex response function (See equations (4) and (5) in reference^5^) which can be directly related to the experimental transfer function H(ω)

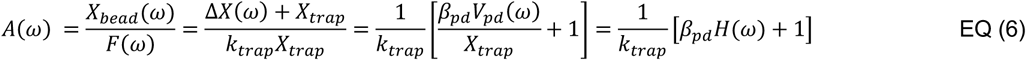

With the previously calibrated k_trap_ and β_pd_ calibrated from EQ (1) and EQ (4) and the experimental H(ω), A(ω) can be solved using EQ (6) and the complex shear modulus G(ω) of the actual cytoplasm can be obtained using EQ (5).

### Simple viscoelastic model fitting

A global fitting algorithm is used to fit simultaneously EQ (1) and EQ (4) to the FRF and power spectrum of the data respectively. As mentioned previously, only frequencies above ∼1 Hz in the FRF and frequencies above ∼200 Hz in the power spectrum are included in the fit (Supplementary Figure 1). Occasionally, despite the addition of an extra pole compensating for the filtering effect observed when data acquisition was done with downsampling (Supplementary Figure 1b, right), or direct sampling where filtering effect was not observed (Supplementary Figure 1b, left), the FRF at higher frequencies showed the presence of an extra pole, causing a shift in magnitude and phase. As the simple viscoelastic model did not include a term to correct for this shift, fitting of the FRF was stopped before reaching these frequencies. The last point in the magnitude plot in Supplementary Figure 1a left shows such a dip in magnitude, indicative of filtering at higher frequencies. Further experiments and quantification of this phenomenon may help to isolate the source and correct the data above ∼500 Hz allowing the higher frequencies to be reliably obtained (frequencies past ∼500 Hz were not included in this manuscript). A weight function was applied to the FRF and power spectrum to account for the large difference in number of points (∼11 for the FRF, and thousands for the power spectrum). A second weight function was applied to the FRF to weight the magnitude response more than the phase as the latter appeared to be more sensitive to sources of error. Each fit begins with an initial guess that is manually found to be close to the solution and with tighter bounds to restrict the algorithm from finding a local minimum in a completely separated region in the parameter space. A second fit is done based on the output of the first fit with looser bounds.

### Bootstrapping and cell mechanics model fitting

Bootstrapping with replacement is done by generating k numbers randomly drawn from 1 to N. N is the total number of data sets per condition, and k is the minimum amount of data available at any given frequency. For example, in the case of untreated WT, the number of data points available per frequency varied from 14 to 17, due to slightly different excitation inputs. In that case, k = 14, and N = 17. This bootstrapping is repeated 1000 times to obtain 1000 bootstrap samples. The mean of these samples is first manually fit to either the power law or dynamic crosslinking model to obtain an initial guess of the parameters. Then, each bootstrap sample is automatically fit to either models using the corresponding initial guess. Each bootstrap sample is fit 3x, starting at the initial guess with stricter bounds, and passing the best parameters found for the next 2 subsequent fits with increasingly looser bounds.

